## Supplemental table 1 for "Functional characterization of a PHF8 processed pseudogene in the mouse genome"

**Supplemental Table 1. Phf8 and Phf8-ps interacting proteins**

| Phf8-specific interacting proteins | Phf8-ps-specific interacting proteins | Phf8 and Phf8-ps common interacting proteins |
|---|---|---|
| Smc2 | Atp1a3 | Polr2a |
| Tdrkh | Rhot2 | Bag1 |
| Tmem165 | Pdpk1 | Abcf2 |
| Atp5h | Scfd1 | Ntpcr |
| Smc4 | Gfpt1 | Dnaja3 |
| Asun | Cul2 | Yme1l1 |
| Smc1a | Fkbp10 | Cbr4 |
| Psme4 | Ndufa10 | Faf2 |
| Edc3 | Qki | Gcn1l1 |
| Sin3a | Abce1 | Slc27a4 |
| Iqsec1 | Ears2 | Phkb |
| Far1 | Pfdn6 | Sucla2 |
| Arcn1 | Pcna | Tubg1 |
| Ints10 | Stub1 | Slc25a12 |
| Hk2 | P4hb | Emd |
| Obfc1 | Pdcl2 | Bag6 |
| Ncapd3 | Tubb4a | Ubxn1 |
| Kif27 | Vbp1 | Sec61a1 |
| Tecr | Btaf1 | Copb1 |
| Tusc3 | Fam78a | Fyn |
| Rad50 | Hars | Araf |
| Mcm5 | Pfdn2 | Slc25a22 |
| Kifap3 | Irak1 | Ythdf2 |
| Psma7 | Nsun2 | Tmem33 |
| Nup188 | Ambra1 | Polr2c |
| Rrp12 | Stip1 | Aifm1 |
| Fanci | Aldh18a1 | Sec22b |
| Psmb2 | Cnp | Dnajc11 |
| Nf1 | Ipo5 | Prkdc |
| Irs4 | Elp3 | Clpb |
| Pomgnt2 | Phkg2 | Wwox |
| Ccdc47 | Praf2 | Xpo1 |
| Stt3a | Sdf4 | Gnai3 |

| | | |
|---|---|---|
| Acad9 | Mad2l1 | Ass1 |
| Atp6v1d | Ipo7 | Dpm1 |
| Psmb1 | Fem1b | Dnajb12 |
| Nsf | Maip1 | Ndufs2 |
| Hyou1 | Pex1 | Afg3l2 |
| Dhcr7 | Nampt | Hsph1 |
| Smc3 | Ndufs3 | Rhot1 |
| Lonp2 | Pfdn5 | Tubb6 |
| Hs2st1 | Galk1 | Slc25a11 |
| Gtf2b | Mlf2 | Vps4a |
| Topbp1 | Dnaaf2 | Ints6 |
| Mark2 | Akap8l | Mcm7 |
| Copb2 | | Arl1 |
| Cby1 | | Cacybp |
| Mogs | | Gnb1 |
| Ctc1 | | Tomm22 |
| Cox15 | | Phgdh |
| Atp2b1 | | Slc25a13 |
| Timm50 | | Ppp6c |
| Fbxw11 | | Lrrc41 |
| Pnpla6 | | Cdipt |
| Simc1 | | Fkbp8 |
| Csnk1g3 | | Pfkl |
| Bcs1l | | Cse1l |
| Chchd3 | | Maged2 |
| Abhd12 | | Huwe1 |
| Pih1d1 | | Glud1 |
| Fam91a1 | | Pla2g6 |
| Sacm1l | | Polr2b |
| Xrn2 | | Copg2 |
| Pigt | | Surf4 |
| Ap3m1 | | Raf1 |
| Tubgcp4 | | Ahsa1 |
| Sgpl1 | | Kdelr2 |
| Srpr | | Wdr11 |
| Smc6 | | |
| Wdr48 | | |
| Ncln | | |
